# An Active Chemo-mechanical Model to Predict Adhesion and Microenvironmental Regulation of 3D Cell Shapes

**DOI:** 10.1101/2020.01.07.897405

**Authors:** Xingyu Chen, Veronika te Boekhorst, Eoin McEvoy, Peter Friedl, Vivek B. Shenoy

**Affiliations:** Center for Engineering Mechanobiology, University of Pennsylvania, Philadelphia, PA 19104; Department of Materials Science and Engineering, University of Pennsylvania, Philadelphia, PA 19104; Department of Genitourinary Medical Oncology, The University of Texas MD Anderson Cancer Center, Houston, TX 77030; Department of Cell Biology, Radboud University Medical Centre, 6525GA Nijmegen, The Netherlands

**Author notes:** Corresponding author: Vivek B. Shenoy, Ph.D., Department of Materials Science and Engineering, University of Pennsylvania, 3231 Walnut Street, LRSM 309, Philadelphia, PA 19104.

**Keywords:** cell, morphology, shape, contractility, adhesions, energies

## Abstract

Cell shapes are known to regulate cytoskeletal organization, stiffness and the ability of cells to migrate and proliferate. Yet a quantitative understanding of the fundamental biochemical and biophysical mechanisms that determine the cell shapes is currently not available. In this study, we developed a chemo-mechanical feedback model to predict how adhesions and the properties of the 3D microenvironment regulate cell shapes. We find that the cells in 3D collagen matrices remain round or adopt an elongated shape depending on the density of active integrins, the level of contractility regulated by mechanosensitive signaling pathways and the density and mechanics of the matrix. While the formation of actin fibers that run along the cell body mediated by integrins and matrix stiffness drive elongation of cells, the cortical and membrane tension resist elongation. Based on the competition between these mechanisms, we derive phase diagrams for cell shape in the space spanned by the density of active adhesions and the level of biochemical signaling that controls contractility. Our predictions are validated by studying the shapes of HT1080 cells cultured in collagen gels of varying densities and using pharmacological treatments to regulate adhesions and contractility. The predictions of the model are found to be in excellent agreement with our experiments and data reported on a number of cell types in the literature.

## I. Introduction

Cells are known to adapt their cytoskeletal structure in response to intracellular signals and changes in the extracellular microenvironment. When cultured on 2D substrates, the cells adopt a more spread and spindle-like morphology with an increase in the substrate stiffness^1^. In 3D, cancer cells are known to adapt to the microenvironment by switching between mesenchymal or ameboid migration strategies while invading into the surrounding matrix^2–5^. Each of these migration modes is characterized by unique cell morphologies. The mesenchymal type cell is generally elongated with strong contractile actin filaments that run along the length of the cells and bridge adhesions, while the ameboid type cells display a round morphology. Emerging evidence shows that the cell shape can have an impact on cell stiffness^6^, contractility^7^, cell fate^8^ and the ability of the cells to proliferate and differentiate^7, 9^. It has been shown that the elevated cytoskeletal tension associated with the spread of cell opens nucleus pores allowing translocation of YAP and TAZ into the nucleus from the cytoplasm, which promotes cell differentiation and suppresses the cell proliferation^10, 11^. Therefore, studying the mechanism of cell shape changes can shed light on the fundamental mechanisms of mechanotransduction.

While cell shapes have been considered in many studies, the mechanisms that regulate cell morphological changes remain poorly understood both qualitatively and quantitatively. In this article, we present a chemo-mechanical approach that integrates signaling with free energy associated with mechanical deformation and active contraction to elucidate the mechanism of cell morphological adaptation to the microenvironment. Specifically, we modeled the signaling pathways that regulate active recruitment of contractile elements in response to the changes in the actomyosin network^12^, the non-linear strain stiffening of the matrix^13^ and the engagement of adhesions to obtain the active free energy of the cell-matrix system. We found that different cell shapes are associated with different levels of biochemical signaling and mechanical stresses and thus have different free energies. We predict the cell shape by minimizing the free energy, and then study how the changes in matrix stiffness, ligand/adhesion density and intracellular Rho-ROCK signaling can impact the cell shape. We validate our theoretical analysis by studying the shapes of HT1080 cells cultured in the collagen matrices as well as data on other cell types reported in the literature. The predictions on the impact of contractility, ECM stiffness and adhesions are tested using pharmacological treatments of cells.

## II. A Chemo-mechanical Model to predict cell shape

We developed a chemo-mechanical model to elucidate the biochemical pathways and physical processes that lead to a switch in the shapes of cells. Our model consists of (Fig. 2a) three components: the cell, the cell-matrix interface, and the matrix. Although many of our results can be generalized, we limit ourselves to the 3D case where we consider an ellipsoidal cell of fixed volume in the matrix and analyze the contributions to the chemo-mechanical free energy of the cell-matrix system (Fig. 1) influences the cell aspect ratio (AR). This free energy includes the cell active potential *F*_*cell*_ that accounts for the active processes that determine the contractility and its polarization, the cell-matrix interfacial energy Γ and the strain energy associated with cell-induced deformation of the collagen matrix *F*_*matrix*_. The total energy of the cell-matrix system *F*_*total*_ is the sum of these three contributions,

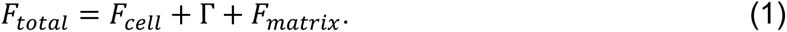

Here we briefly describe the key aspects of these components; more details are included in the supplementary information.

**Fig. 1.**
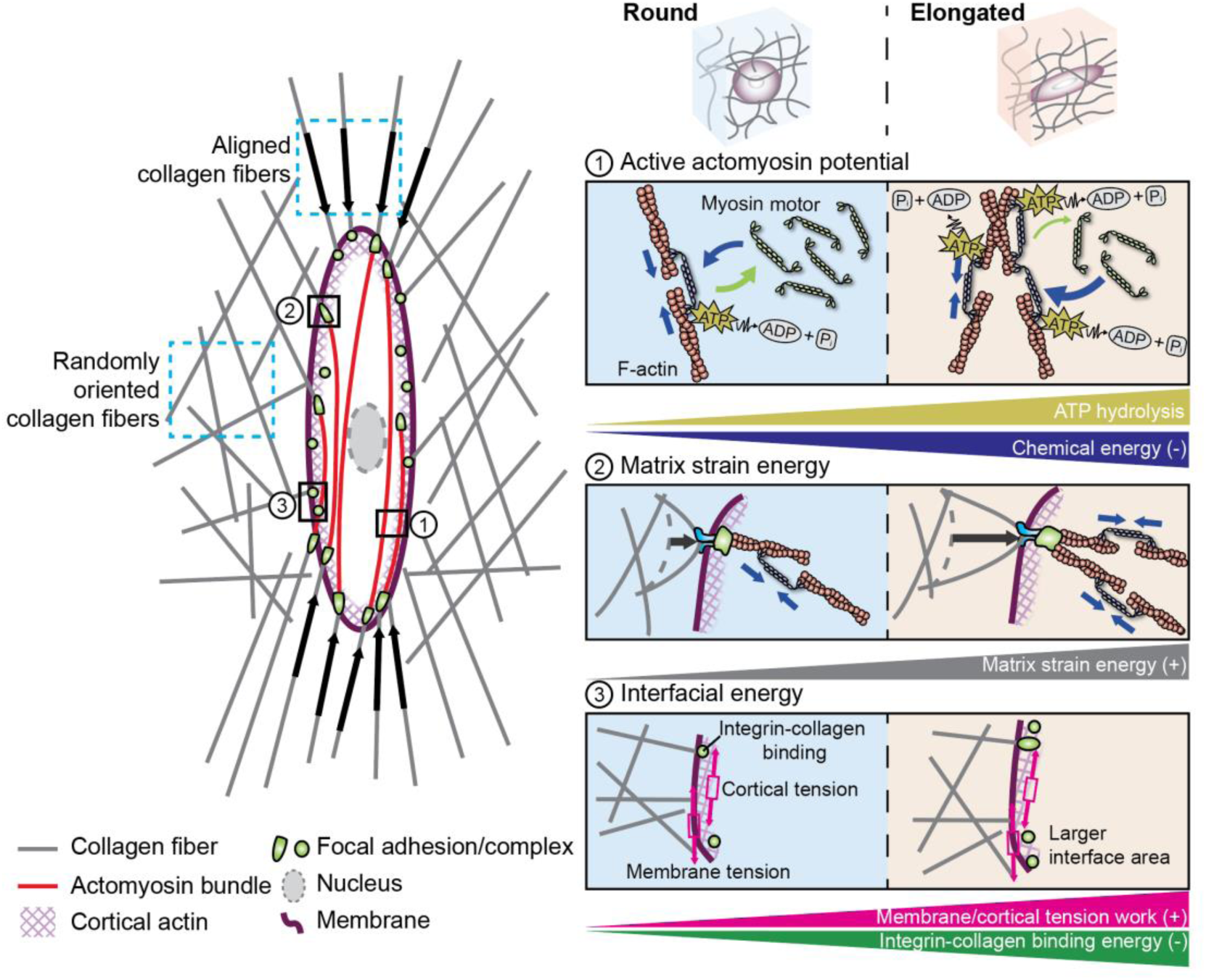
This model considers the cell active potential *F*_*cell*_, the matrix strain energy *F*_*matrix*_ and the interfacial energy Γ as marked in the schematic. As the cell elongates, myosin motors are recruited and polarized along the cell long axis, forming aligned contractile actin filaments. This actomyosin configuration has a lower chemical energy, leading to a lowered *F*_*cell*_. In the matrix, the larger deformation induced by the contraction of elongated cell increases the matrix strain energy *F*_*matrix*_. Lastly, the binding affinity between the integrins and collagen favors a larger cell-matrix interface area associated with elongated shapes, while the cortical and membrane tension work against it. The competition among the individual energy components determines the shape of the cell.

**Fig. 2.**
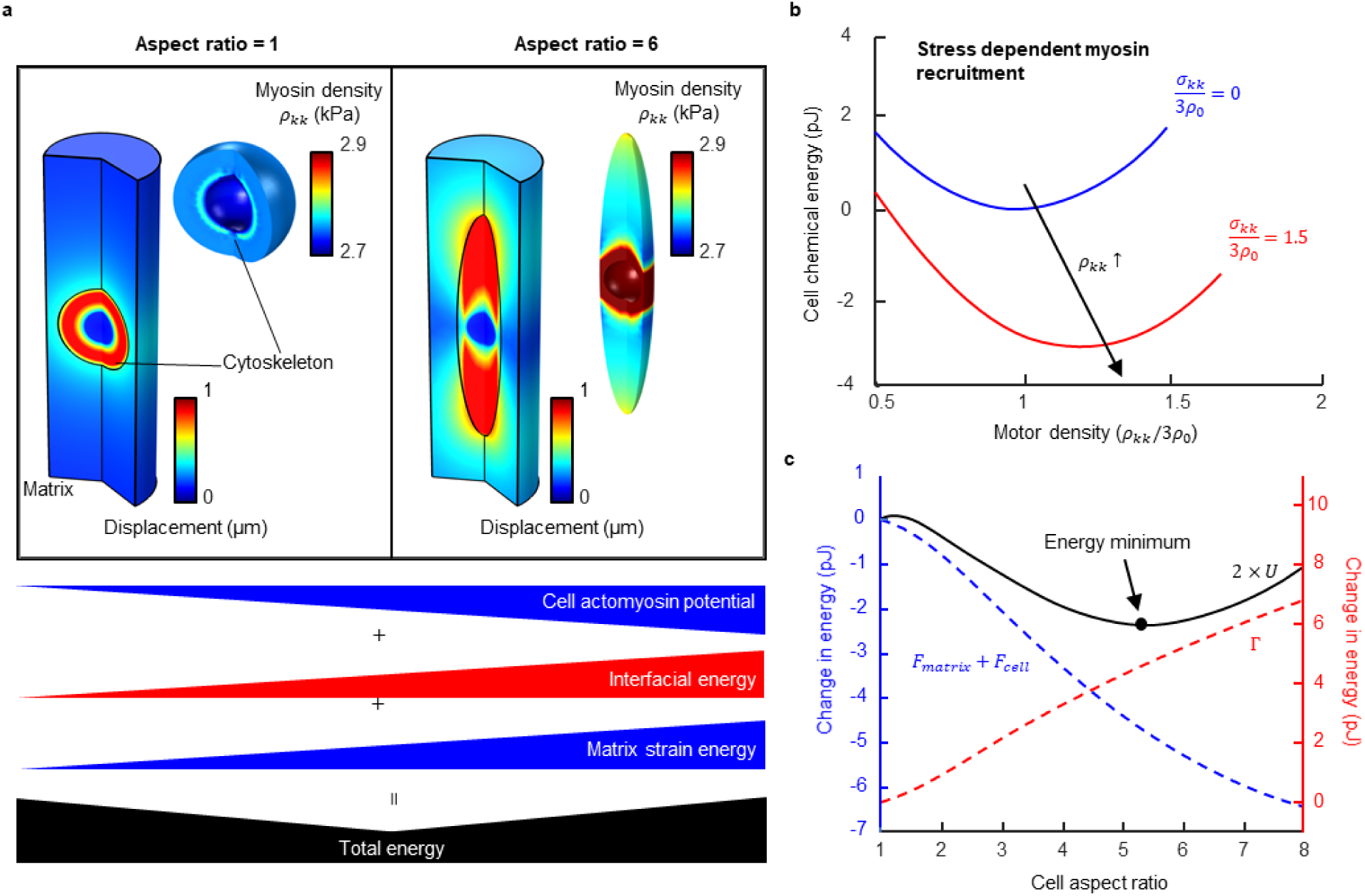
(a) The displacement field induced by cell contraction, and the spatial variation of the density of myosin motors for the case of cells with aspect ratios of 1 and 6. (b) The change in cell chemical energy. (c) The changes in different contributions to the energy as the cell elongates. The competition of the cell active potential, matrix strain energy and interfacial energy leads to a non-monotonic total energy with respect to the aspect ratio of the cell. The preferred aspect ratio corresponds to the case when the overall free energy is the lowest.

### Cell active potential

The cell potential *F*_*cell*_ represents the energy involved in the assembly of actomyosin networks and the generation of contractile forces. Microscopically, myosin motors can convert chemical energy from ATP hydrolysis into mechanical work. The active forces of myosins lead to the contraction of the cell, deforming the cytoskeleton, nucleus, and transmission of forces to the surrounding matrix through adhesions. The cell a potential *F*_*cell*_ contains a passive part *F*_*passive*_ and an active part *F*_*active*_, *F*_*cell*_ = *F*_*passive*_ + *F*_*active*_, which we describe next.

The passive part describes the strain energy stored in the cell nucleus and the cytoskeleton when deformed by contractile forces.

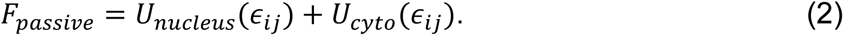

Here, *U*_*nucleus*_ and *U*_*cyto*_, which are functions of the strain tensor *ϵ*_*ij*_, denote the strain energy stored in the nucleus and cytoskeleton, respectively (Refer to SI for details).

The active part *F*_*active*_ describes how mechanical stresses can lead to both recruitment and polarization of actomyosin networks. In the absence of stresses, the contractility is isotopic and small, while in the presence of stress, the contractility of cells increases, and they polarize along the axes of the largest tensile stress. We use the tensor *ρ*_*ij*_ to denote the density of myosin motors, treated as force dipoles with the indices i and j denoting the components of force and the direction along which these forces are offset, respectively

(Refer to Fig. S1). The active free energy *F*_*active*_ can be written in terms of this contractility tensor as

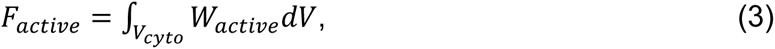

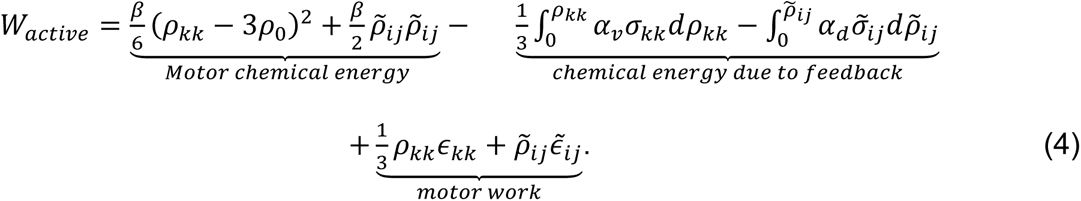

Table 1 lists the physical interpretation of each term in this equation.

**Table 1.** The physical meaning of each term in Eq. 5.

| Terms in Eq. 4 | Physical meaning |
| --- | --- |
| Motor chemical energy | The chemical energy associated with the myosin binding and unbinding process in the absence of stresses. |
| Feedback | Impact of external stress on the recruitment of contractile elements through force-activated signaling pathways (Rho-ROCK, $\text{Ca}^{2+}$). |
| Motor work | Motor mechanical work converted from chemical energy |

Here *V*_*cyto*_ denotes the volume of the cytoplasm. *ρ*_*kk*_ is the total density myosin motors, and the deviatoric part *ρ̃*_*ij*_ characterizes the polarization of the contractility tensor. The first two terms representing the chemical energy ensures that the contractility is isotropic (*ρ̃*_*ij*_ = 0) in the absence of stresses transmitted to the cytoskeleton from the ECM. The third and fourth terms account for the recruitment and polarization of myosin motors in response to stresses transmitted to the adhesions. The feedback parameters *α*_*v*_ and *α*_*d*,_ that describe the mechanosensitive feedback pathways (such as the Ca^2+^ and Rho-ROCK signaling), relate the total density myosin motors *ρ*_*kk*_ to the mean stress *σ*_*kk*_ and the polarization of contractility *ρ̃*_*ij*_ to the anisotropy in the state of stress *σ̃*_*ij*_ (Eq. S3). The last two terms represent the mechanical work done by the motors. A key feature of the active free energy is that it accounts for the phenomenon of durotaxis, where the chemical free energy is lower if the cell is in a stiffer environment and is able to spontaneously increase its contractility. Similarly, as we show below, when cells are elongated, stresses are anisotropic and the contractility is polarized along the long axis resulting in a reduction in the free energy (which becomes more negative) compared to spherical cells with isotropic contractility as shown in Fig. 2c.

### Interfacial energy

The interfacial energy Γ characterizes the energy of the cell-matrix interface, including the contributions from the adhesions, the actin cortex, and the membrane. At the cell-matrix interface, the receptors (such as integrin) on the cell surface spontaneously bind to collagen ligands forming adhesions and lowering the energy. Since cell elongation increases cell surface area and can expose more collagen ligands available for binding, which facilitate the formation of adhesions and thus lower the interfacial energy Γ. Here we assume that the number of adhesive receptors per unit area (receptor density *d*_*re*_) of the cell is fixed. When the receptor density is high, it becomes favorable for the cell to adopt an elongated shape to maximize the number of bonds between the cell and the matrix. On the other hand, the membrane and cortical tension work against the increase in surface area, and therefore do not favor elongation of the cell. We captured these features by writing the interfacial energy Γ as

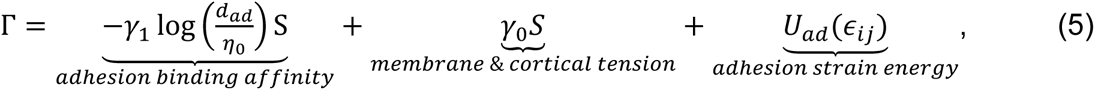

where *S* denotes the area of the interface, *γ*_0_ denotes the contribution from the membrane and cortical tension. The parameter *γ*_1_ is the change of energy associated with the formation of each receptor-ligand bond, and *d*_*ad*_ denotes the density of adhesions, which can be calculated by

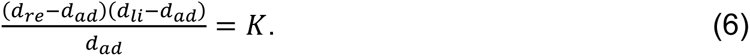

Here *d*_*re*_ and *d*_*ligand*_ denote the density of receptors and ligands respectively. K denotes the equilibrium constant of the receptor-ligand binding reaction. In Eq. 5, we also include the strain energy stored in the adhesions *U*_*ad*_(*ϵ*_*ij*_) which are deformed as they transmit forces from the cytoskeleton to the ECM (refer to SI for details).

### Matrix strain energy

The matrix strain energy *F*_*matrix*_ characterizes the strain energy stored in the matrix when deformed by the cell. It is calculated by integrating the strain energy density (strain energy per unit volume) *W*_*matrix*_ over the extracellular volume, *V*_*matrix*_

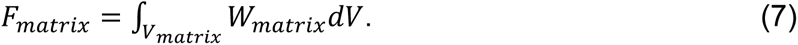

When the collagen network deforms, a fraction of the initially randomly oriented fibers align along the directions of tensile principal strains. The increased alignment strain stiffens the network in an anisotropic manner in that direction. The strain energy density function *W*_*matrix*_ consists of the contributions from both the unaligned fibers and aligned fibers, which can be written as

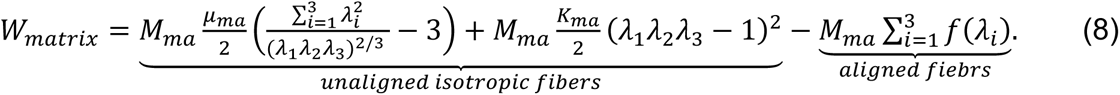

Here *μ*_*ma*_ and *K*_*ma*_ are material properties characterizing the shear and bulk response of the matrix and *λ*_*i*_ (*i* = 1,2,3) denote the principal stretches. The power-law function *f* (see SI for details) captures the matrix stiffening in the tensile principal directions. The scale factor *M*_*ma*_ is introduced to correlate the collagen concentration with stiffness.

## III. Results

### The shape of the cell is determined by the competition between the active potential and the membrane and matrix deformation energy

To predict cell shape, we first investigated the shape-dependence of the free energy components, including the cell active potential *F*_*cell*_, matrix strain energy *F*_*matrix*_ and the interfacial energy, Γ. The extreme cases of a round (AR = 1) cell and an elongated (AR =6) cell are considered in Fig. 2. As the cell active potential *F*_*cell*_ is determined by the cell stress and contractility, we first explored how these quantities changed with cell shape. Both the cell volumetric stress (*σ*_*kk*_) and the deviatoric stress along the cell long axis (*σ̃*_33_) were higher in the case of the elongated cells (Fig. S2a). In response to elevated stress, the cell recruits additional phosphorylated myosin motors through force-activated signaling pathways (e.g., Rho-ROCK, Ca^2+^) and polarizes to form aligned contractile actin filaments, as indicated by the high levels of the mean contractility (*ρ*_*kk*_) and its polarization along the long axis (*ρ̃*_33_) (Fig. 2a & S2b). With increased myosin recruitment and polarization (Fig. S5), the cell active potential *F*_*cell*_ becomes more negative as the cell elongates (Eq. 5). In the matrix, while a radially symmetric displacement field was predicted surrounding the round cell, the elongated cell most significantly deformed the matrix proximal to the cell tips due to its polarized contractility. The magnitude of the displacement decayed as the distance from the cell increased, in good agreement with experimental observations^14, 15^ of the displacement field around a single cell in 3D collagen matrices. As highly elongated cells are shown to significantly deform the matrix, there is an associated increase in the strain energy, *F*_*matrix*_. This acts as an energetic penalty that prevents the cell from taking on extremely large aspect ratios. Finally, the increase in interfacial energy Γ shown in Fig. 2c is attributed to two factors: (i) the adhesion strain energy (Eq. 6) increases with cell elongation due to high levels of contractility stretching the bonds between the cell and the matrix, and (ii) the cell surface area scales both the integrin-collagen binding energy and the membrane tension (Eq. 6). The surface area is lowest when the cell assumes a spherical shape (given a fixed volume). As a result, the overall interfacial energy increases with increasing cell aspect ratio as shown in Fig. 2c, acting as a further penalty to cell elongation.

Thus with increasing cell aspect ratio, the cell active potential decreases, while both the matrix energy and the cell-matrix interfacial energy increase (Fig. 2c). This competition leads to a non-monotonic profile of the overall cell free energy with respect to its shape. Increasing the cell aspect ratio initially leads to a lowering of the free energy, and a minimum free energy configuration can be identified which corresponds to the optimum cell aspect ratio of (∼5.2 in Fig. 2c). Beyond this optimum value, the interfacial energy dominates the competition, leading to an increased cell-matrix free energy for highly elongated cells. With the understanding of how variations in cell shape affect the cell-matrix system free energy, we next explore the influence of matrix stiffness, myosin activity, and adhesions on cell elongation.

### Cell elongation in stiff matrices is driven by increased actomyosin activity

A range of cell aspect ratios were analyzed in matrices of increasing stiffness, and the cell-matrix system energy *F*_*total*_ for was computed for each of the cases. When the cell is embedded in a soft matrix (matrix stiffness scaling factor *M*_*ma*_ = 0.8), cell contractility is not significant (Fig. 2b) and the change in cell active potential *F*_*cell*_ (with increasing cell aspect ratio) is low. The system free energy is dominated by the interfacial energy Γ, which monotonically increases with the cell aspect ratio. In this case, the minimum free energy configuration corresponds to a round cell shape (i.e. with aspect ratio = 1), as any attempt by the cell to change its shape leads to an increase in the cell-matrix system (see Fig. 3c).

**Fig. 3.**
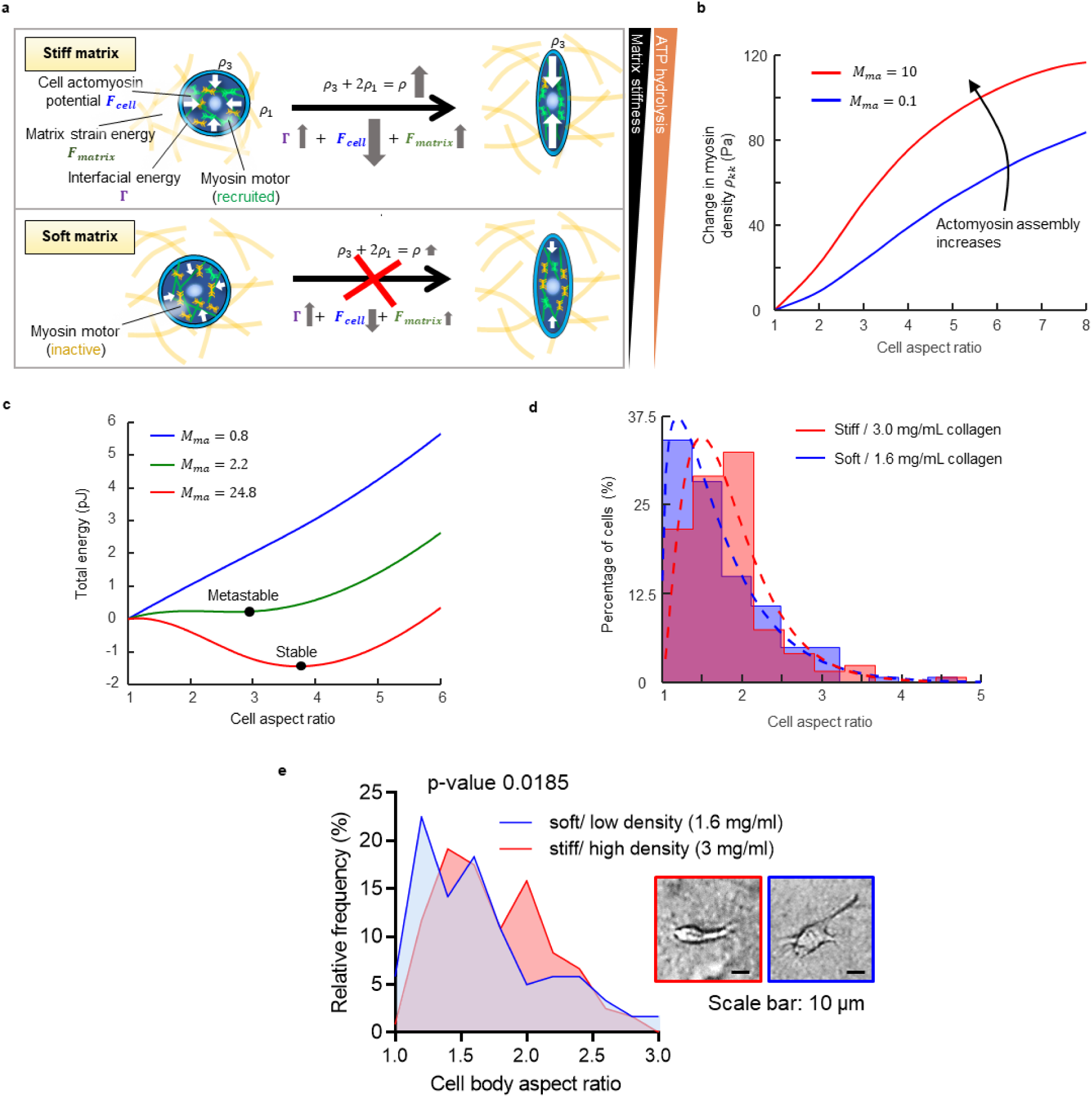
(a) Schematic showing the impact of matrix stiffness on cell elongation. In a stiff matrix, elongation is associated larger increase in the cell volumetric stress and actomyosin assembly. The reduction in the actomyosin active potential reduction favors elongation in stiffer matrices compared to the softer matrices. (b) The relative myosin density as the cell elongates in stiff or soft matrices. (c) The total cell-matrix system energy when the cell adopts different shapes. As the matrix stiffness increases, the cell adopts an elongated shape with lower overall free energy. The (d) *in silico* and (e) *in vitro* histogram of aspect ratio of cells cultured in 1.6 or 3.0 mg/mL collagen matrices (curves are fitted using Gamma distribution).

In a stiff matrix, our model predicted that cells become more contractile, with increased myosin recruitment and polarization shown by elevated levels of the mean contractility *ρ*_*kk*_ and its polarization along the long axis *ρ̃*_33_, respectively (Fig. 3b). This prediction is supported by experimental observations of fibroblasts adhered to flexible cantilevers^16^, where they force exert increases with the stiffness of the cantilever. Furthermore, the increase in the volumetric and deviatoric part of the contractility tensor *ρ*_*kk*_ and *ρ̃*_33_ (associated with increasing cell aspect ratio) is more pronounced on stiffer matrices. As discussed previously, the cell active potential becomes more negative with increasing myosin recruitment and polarization (via Eq. 5). Thus, when the matrix stiffness is increased (*M*_*ma*_ = 2.2), the contribution of the cell active potential becomes more significant. The monotonic trend gives way to a more evident competition between the active potential and the interfacial and matrix contributions, with a minimum in free energy predicted for an aspect ratio of ∼2.9. When the matrix stiffness is further increased (*M*_*ma*_ = 24.8), the cell-matrix system energy is dominated by the cell active potential up to an aspect ratio of ∼3.8, i.e. the minimum free energy state. Note that in the case of intermediate stiffness (*M*_*ma*_ = 2.2) there existed two local energy minima at aspect ratios of 1 and 2.9, while only one energy minimum was observed for high stiffness (*M*_*ma*_ = 24.8), which suggests that elongated shapes are more stable in stiffer matrices.

To test our model predictions, we measured the aspect ratios of HT1080 cells cultured in 1.6mg/mL or 3mg/mL collagen matrices. In agreement with our model, the cells in the 3mg/mL matrix (higher stiffness) exhibited significantly higher ARs relative to cells in the 1.6mg/mL matrix (Fig. 3d & Fig. S5). Similar results have also been observed for melanoma cells cultured in collagen from 0.1 mg/mL to 2mg/mL^17^. The elongated cells we also found to show significantly higher actomyosin recruitment and polarization, consistent with our model predictions.

### Cells become round when the actomyosin activity is disrupted

Having determined that actomyosin activity is essential to facilitate cell elongation, we next investigated the role of mechanosensitive signaling (e. g. Rho-ROCK, Ca^2+^ pathways) on the regulation of cell shape (Fig. 4a) by varying the feedback strength *α* (*α*_*v*_ = *α*_*d*_ = *α*) and chemical stiffness *β*. When the cell-matrix system reaches a steady state (see SI for details), the cell contractility *ρ*_*ij*_ was found to be correlated with stress via these two parameters (Eq. S1). In the previous sections of this paper, we have demonstrated that actomyosin activity favor elongated cell shapes and that the cell assumes such elongated states if the cell active potential *F*_*cell*_ is comparable in magnitude to other passive contributions. Therefore, disrupting this active feedback will lead to round cell shapes. Commonly used disruptive treatments include Y-27632 (downregulates ROCK expression) and blebbistatin (inhibits myosin binding) (Fig. 4a). These treatments were simulated by lowering *α* in our model, which reduces the dependence of the contractility tensor *ρ*_*ij*_ on force signaling (via Eq. S1). We observed that, as the cell elongated, the changes in *ρ*_*kk*_ (Fig. 4b) and *ρ̃*_33_ were lower when the feedback was inhibited (i.e., *α*/*β* reduced from 0.85 to 0.7). As a result, the contribution of the cell active potential became less significant (Eq. 5), driving the cell to adopt more rounded configurations. The predicted cell shapes with different levels of feedback strength, *α* are shown in Fig. 4c. Decreasing *α* reduces cell elongation, and at a critical value, the cell shape abruptly becomes round and continues to maintain this shape for smaller values of the mechanochemical feedback parameter.

**Fig. 4.**
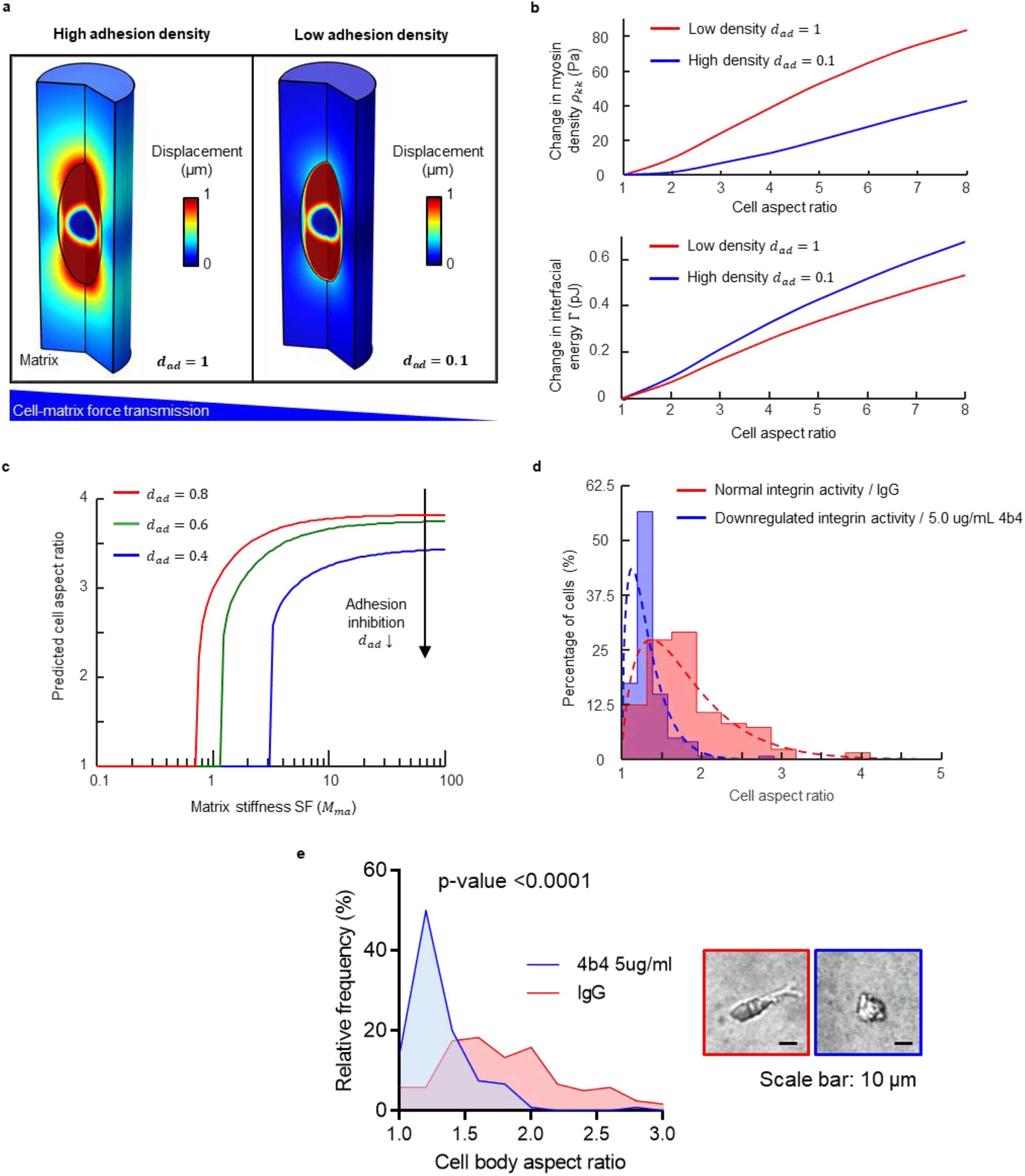
(a) Schematic showing the impact of mechanosensitive feedback or myosin inhibition on cell shapes. (b) The relative myosin density as the cell elongates (*d*_*ad*_ = 1, *M*_*ma*_ = 0.1). (c) The predicted cell aspect ratio with different feedback strength (*α*/*β*), adhesion density (*d*_*ad*_), and matrix stiffness (*M*_*ma*_). The (d) *in silico* and (e) *in vitro* histogram of aspect ratio of cells with or without treatment of blebbistatin that inhibits he activity of myosin (curves are fitted using Gamma distribution).

Our model predictions were validated by measuring the ARs of HT1080 cells treated with blebbistatin. Consistent with the predictions of our model, the ARs of treated cells were significantly lower than those of the controls (Fig. 4d & Fig. S5). The same trend has also been observed for fibroblasts where the cells exhibited significantly rounder shapes when treated with blebbistatin. In addition, previous studies have shown that Y-27632 (ROCK inhibitor) can reduce cell elongation^19^, further confirming the validity of this model. It should be noted that when we treated HT1080 cells with blebbistatin, some of the cells became dendritic with long protrusions originating from various locations on the cell surface. In this case to obtain an unbiased estimate of the aspect ratio, we fit an ellipse (Fig. S6) with the maximum area to the inner cell boundary (as proposed by Thievessen et al.^15^). This measurement of the shape of the cell body (excluding protrusions) gave excellent agreement with model predictions.

In our experiments, we observed HT1080 cells became slightly elongated when treated with low doses blebbistatin, contrary to model predictions. This deviation may be attributed to the impact of blebbistatin on cell cortical tension. While our current computational model assumes that cortical tension (*γ*_0_) is constant, it can be reduced upon treatment with blebbistatin^20^. Therefore, blebbistatin can also regulate the cell shape by decreasing Γ (in addition to *F*_*cell*_ as discussed in previous sections). Such an interaction between cortical contractility and the system free energy influences cell shape in a non-linear fashion. At low blebbistatin concentrations, the cell elongated due to the decreased cortical tension. Only when the blebbistatin concentration reached a critical level, the cell started to become less elongated due to the decreased actomyosin activity.

### Disrupting integrin-based adhesions downregulates actomyosin activity and prevents cells from elongating

Finally, to analyze the role that integrins play in the regulation of cell shape, the variation the overall free energy was studied by varying cell aspect ratios in the case of high (*d*_*ad*_ = 1) and low (*d*_*ad*_ = 0.1) adhesion densities. Here the density of adhesions, *d*_*ad*_ is correlated with both the density of receptors and ligands through Eq. 6. Simulations reveal that a cell will typically assume a more rounded shape when adhesions are impaired (i.e. *d*_*ad*_ = 0.1). With reduced adhesion density, the matrix displacements were lower than that computed with a high ligand density (Fig. 5a). As the adhesions mechanically link the cytoskeleton to the matrix, a reduced density impairs cell-matrix force transmission (i.e. the cell cannot effectively contract the matrix). Within the cell, both the volumetric and deviatoric stresses decreased (Fig. S4a), and the behavior resembled that of a cell in soft matrix. Thus, inhibition of integrins has a similar effect as a reduction in matrix stiffness, i.e. downregulation of actomyosin activity (described in detail in the previous section). Our model predicted that the mean contractility *ρ*_*kk*_ (Fig. 5b) and its polarization along the long axis *ρ̃*_33_ (Fig. S4b) decreased with the adhesion density, consistent with previous studies^21^. Furthermore, when the cell elongates, the surface area increases and exposes more ligands in the collagen to the integrins, facilitating the formation of adhesions and lowering the interfacial energy Γ. When the density of receptors on the cell surface decreases, a smaller number of new adhesions can form as the cell elongates. The increase of interfacial energy Γ becomes more pronounced and penalizes the elongation of the cell. In Fig. 5c, we show the coupled effect of adhesion density and matrix stiffness on cell shape. For any elongated cell shape, our model predicted that a low bond density causes the cell to become more rounded.

**Fig. 5.**
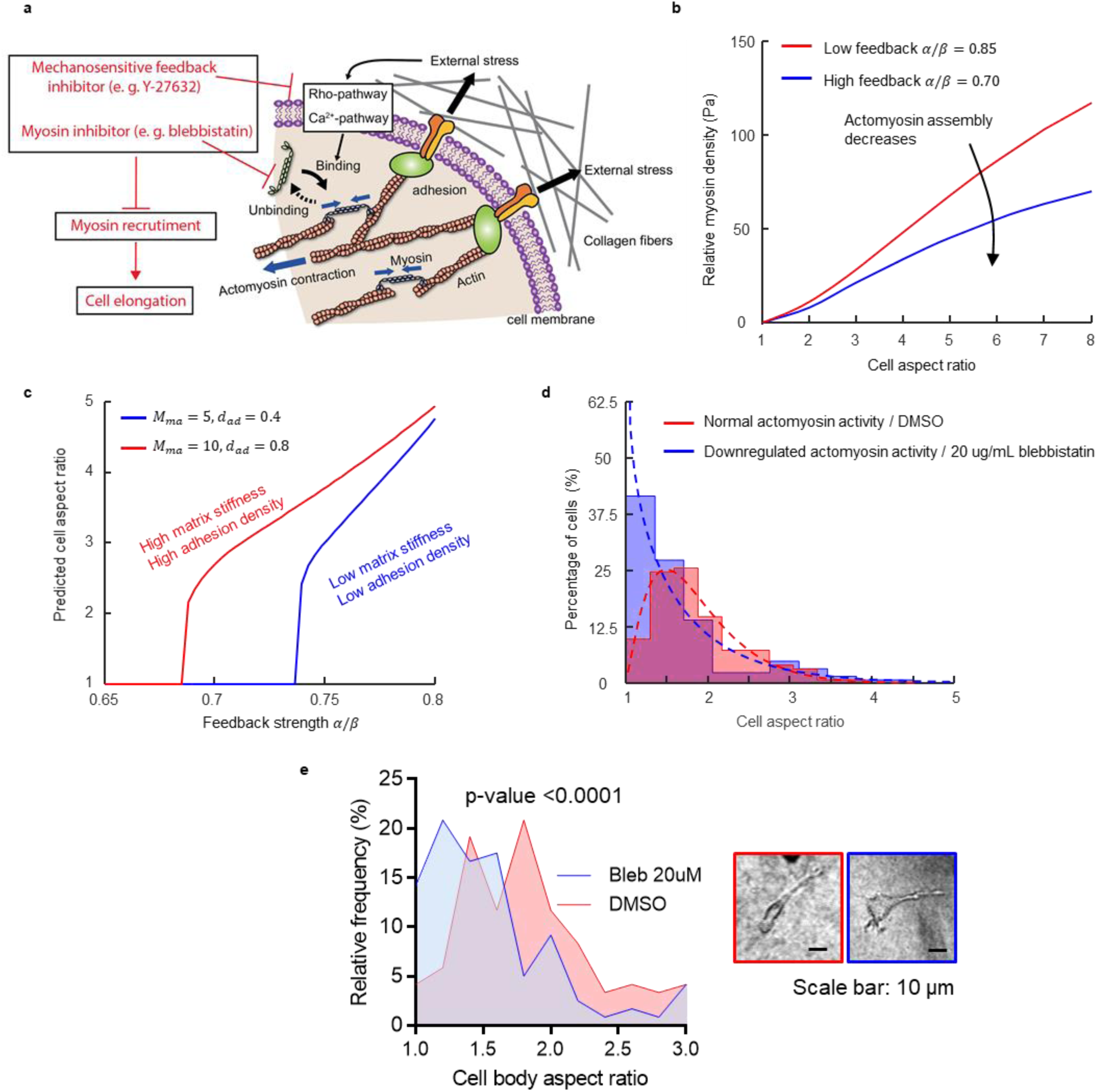
(a) The displacement field in the matrix and the myosin density the cell for low (d_ad_ = 0.1) and high adhesion densities (*d*_*ad*_ = 1). (b) The impact of adhesion density on the actomyosin assembly and the interfacial energy. (c) Cell aspect ratios predictions for different adhesion densities *d*_*ad*_ and matrix stiffness *M*_*ma*_. The (d) *in silico* and (e) *in vitro* histogram of aspect ratio of cells treated with or without 4B4 mAb that downregulates the activity of integrins (curves are fitted using Gamma distribution).

Our predictions are confirmed by experiments where HT1080 cells are treated with 4B4 mAb, which leads to downregulation of β1-integrins^22^. With fewer available receptors on the cell surface, the adhesion formation is impaired, and as captured by Eq. 6, the adhesion density *d*_*ad*_ decreases. Consistent with our model prediction, cells treated with 5µg/mL 4B4 mAb exhibited significantly lower cell AR (Fig. 5d & Fig. S5). Our conclusions also agree with previous studies. Wolf et al.^23^ demonstrated the loss of elongation and polarization in HT1080 cells treated with 4B4 β1-integrin inhibitor; Cukierman et al.^24^ reported that the fibroblast elongation reduced when cells were treated with *α*_*v*_-integrin inhibitor; Castelló-Cros et al. observed that the aspect ratio of MDA-MB-231 cells decreased under the treatment of β1-integrin inhibitor mAb13^25^.

### Predicted cell shape phase diagram

In order to illustrate the integrative impact of matrix stiffness *M*_*ma*_, adhesion density *d*_*ad*_ and feedback strength *α*, we have constructed a cell shape phase diagram. Two slices of the 3D phase diagrams are plotted (Fig. 6) as a function of feedback strength *α* and adhesion density *d*_*ad*_ for the cases of soft (*M*_*ma*_ = 1) and stiff (*M*_*ma*_ = 10) matrices. As shown in Fig.6, in both cases the cells with a high adhesion density and feedback strength adopt elongated shapes. Consistent with this prediction, when we treated HT1080 cells with either integrin-inhibiting mAb or blebbistatin (myosin inhibition), we observed that cells became significantly less elongated (Fig. 4d). In addition, our model also predicted that cells become more elongated in stiffer matrices. For identical parameters (e. g. *α*/*β* = 0.7, *d*_*ad*_/*η*_0_ = 0.8), cells that are rounded in soft matrix become elongated in the stiff matrix. In experiments, we compared the aspect ratio of the HT1080 cells cultured in matrices of low (1.6 mg/ml) and high (3.0 mg/ml) collagen concentrations and found that cells in a matrix with higher collagen concentration show significantly more elongated shapes (p < 0.02).

**Fig. 6.**
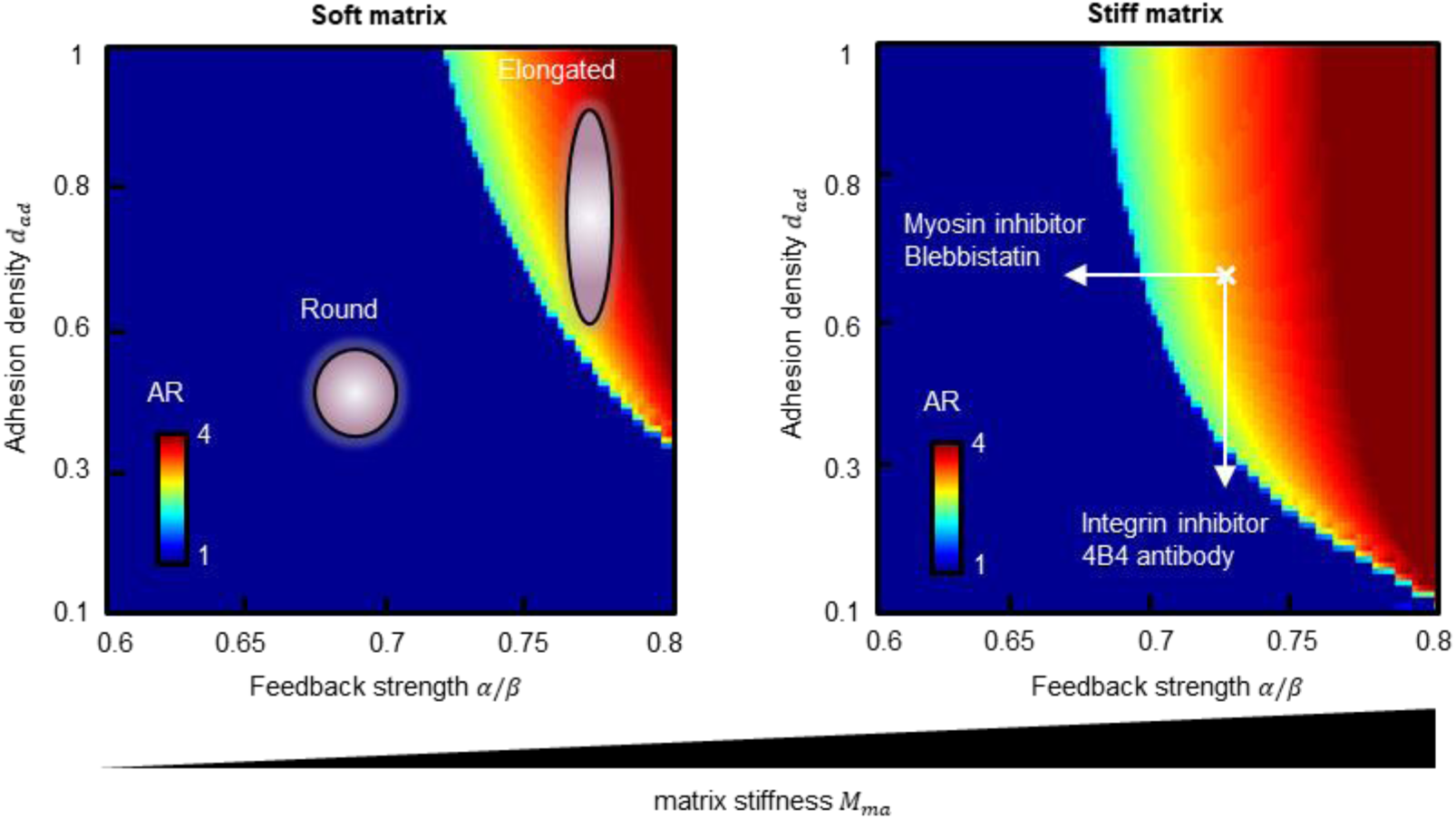
The predicted phase diagram of cell aspect ratio in soft (*M*_*ma*_ = 1) and stiff (*M*_*ma*_ = 10) matrix.

## IV. Discussion

In this study, we have developed a computational model that calculates the free energy of cell embedded in matrix and have elucidated the driving forces underlying cell elongation. We have successfully predicted the dependence of cell shape on matrix stiffness, adhesion density, and mechanosensitive feedback mechanisms. Specifically, we considered how the cell shape emerges as a consequence of the competition between the cell active potential, the cell-matrix interfacial energy, and the matrix strain energy. We demonstrate that the cell active potential decreases as the cell elongates, indicating the actomyosin contractility associated with the aligned actin fibers that run along the cell body. In contrast, the interfacial energy and the matrix strain energy increase with the cell aspect ratio, penalizing cell shapes with a high aspect ratio. The competition among these contributions allowed us to identify the shapes cells are most likely to adopt depending on the properties of the matrix, the density of integrin and the strength of mechano-chemical signaling. The model was validated by measuring the shapes of HT1080 cells cultured collagen matrices and by studying the impact pharmacological treatments on cell shapes. In agreement with our model, the HT1080 cells were significantly less elongated when treated with blebbistatin or 4B4 mAb, or a combination of both. Such results also agree well with previous experiments using fibroblasts^15^ and breast cancer cells^25^, indicating that the mechanisms of shape change elucidated by our model are fundamental across different cell types.

While we limited ourselves to analyzing the impact of matrix stiffness, receptor density, and force-activated signaling, our model can be applied to a variety of other systems. For example, we have identified that cells adopt round shapes when the force transmission between the cell and matrix is impaired. This conclusion also applies to the impact of vinculin, which strengthens the cell adhesions and promotes traction force generation. Thievessen et al.^15^ demonstrated that vinculin-knockout fibroblasts exerted much less contractile force on the surrounding matrix. Indeed, with significantly decreased cell-matrix force transmission, these cells exhibited much lower elongation and polarization^15^. In addition, our model also explicitly considers the collagen ligand density (*d*_*li*_). As captured by Eq. 6, fewer cell-matrix adhesions *d*_*ad*_ can form with a reduction of ligand density. Thus, similar to the impact of reducing the receptor density *d*_*re*_, cells tend to adopt less elongated shapes. In agreement with this conclusion, previous experimental studies aimed to alter ligand density independent of the matrix elasticity have shown that decreasing ligand density inhibits cell elongation^26^.

It is important to note that we consider the energy involved in the actomyosin network that are firmly connected to the matrix through integrins, and thus the model predictions are for the cell body without protrusions. As proposed by Thievessen et al.^19^, when we measured the shapes of HT1080 cells to validate the model, we fit an ellipse inside the cell and found excellent agreement with our model predictions. While the impact of protrusions does not invalidate our shape analysis, it is a critical factor in the cell shape change^25^ and worthy of further exploration (more details in SI). When the round cell contracts in the initially random matrix, the actomyosin activity of the cell increases and causes deformation in the matrix. As we have discussed, the cell will become polarized and elongated driven by the energy minimization. We have shown that the cell-induced matrix deformation becomes maximal at the two ends of the cell long axis (Fig. 2). This aligns the matrix fibers in the direction along the cell long axis, which provides contact guidance for the formation of protrusions. Therefore, the cell with an elongated body tends to form protrusions at the two ends, making the whole cell even more elongated. (Fig. 7) In summary, our chemo-mechanical model and experiments elucidated the underlying competitions among different energy components which dictate the change of cell shape. In addition to providing an explanation for a variety of experimental observations, this model also serves as a theoretical framework to study the shape change of cells under different conditions.

**Fig. 7.**
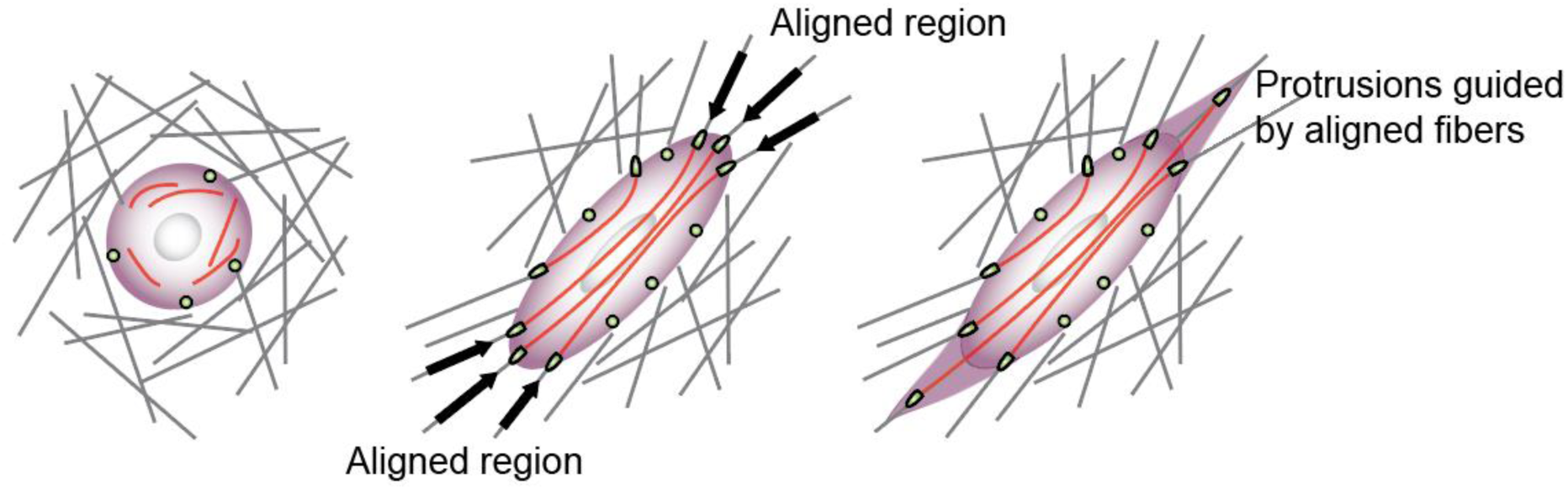
As the round cell contract in the initially randomly aligned matrix, the cell becomes polarized and elongated driven by the energy minimization. The elongated cell deforms the matrix the most at the ends of the cell, aligning the matrix fibers in those regions along the direction of the cell long axis. The aligned fibers provide contact guidance for the formation of protrusions, making the whole cell more elongated.

---

**Fig. S1.**
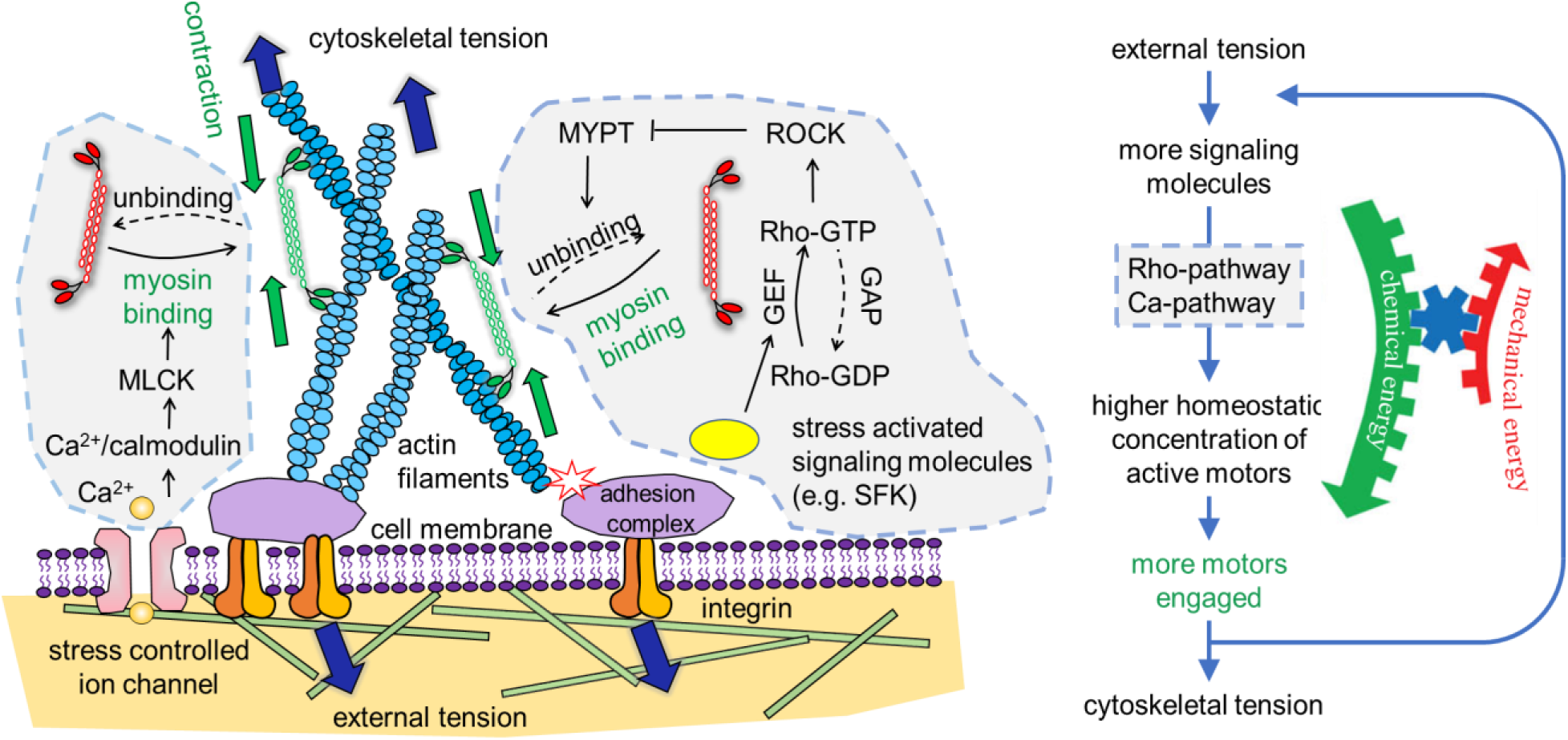
Schematic description of the cell contractility model. The Rho-ROCK and Ca^2+^-pathways are regulated by the stresses from the ECM. When the cell is under tension, a series of biochemical process is activated, and ultimately, more myosin motors switch from inactive state (red) to active state (green) binding to the cytoskeleton, creating more force dipoles in the cell. Collectively, these force dipoles lead to contraction of the ECM.

**Fig. S2.**
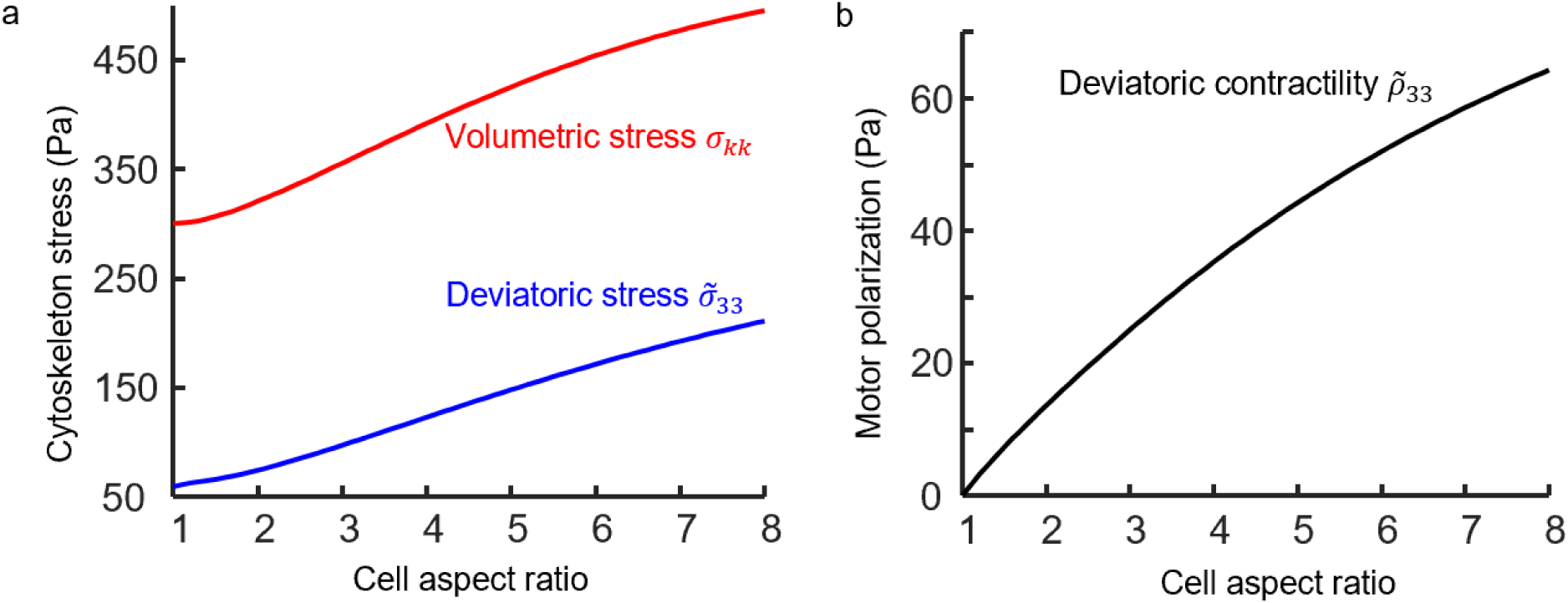
(a) The volumetric and deviatoric stress in the cytoskeleton. (b) The deviatoric contractility tensor in the cytoskeleton.

**Fig. S3.**
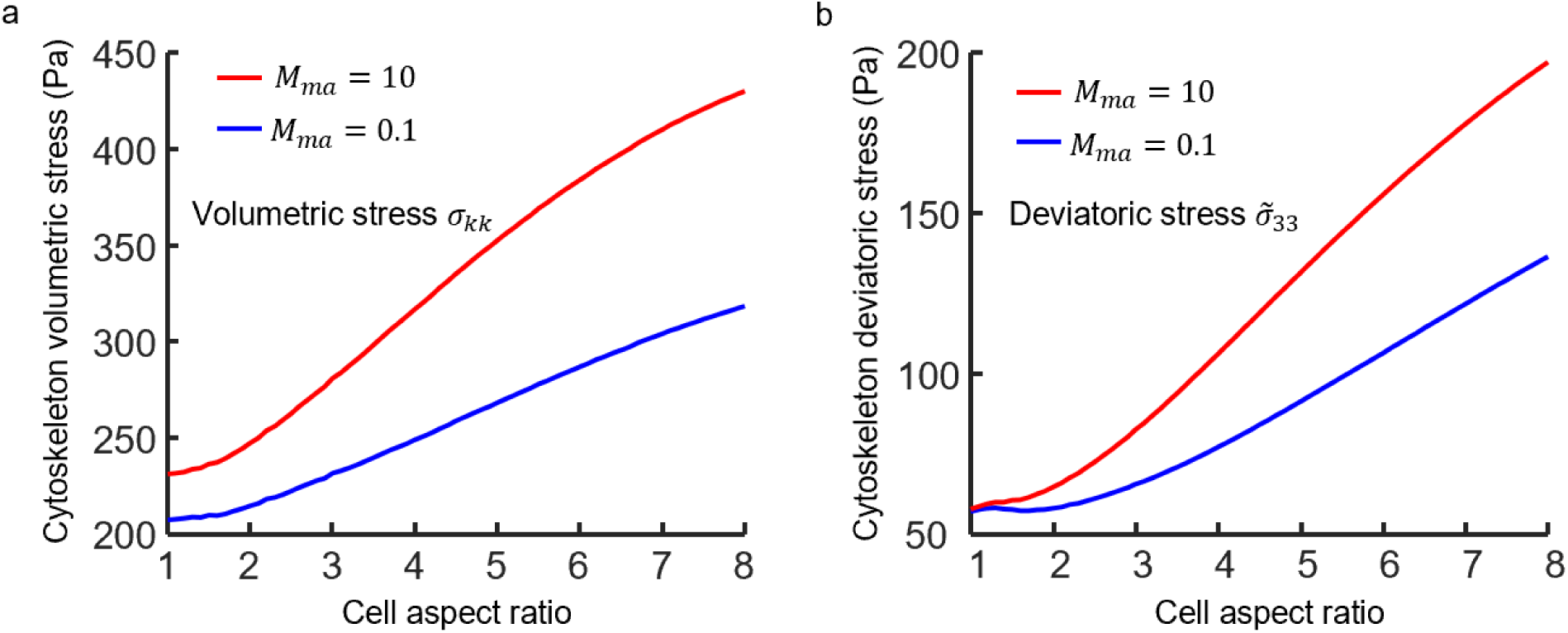
The cytoskeleton (a) volumetric stress and (b) deviatoric stress when the cell is in stiff (*M*_*ma*_ = 10) or soft (*M*_*ma*_ = 0.1) matrix.

**Fig. S4.**
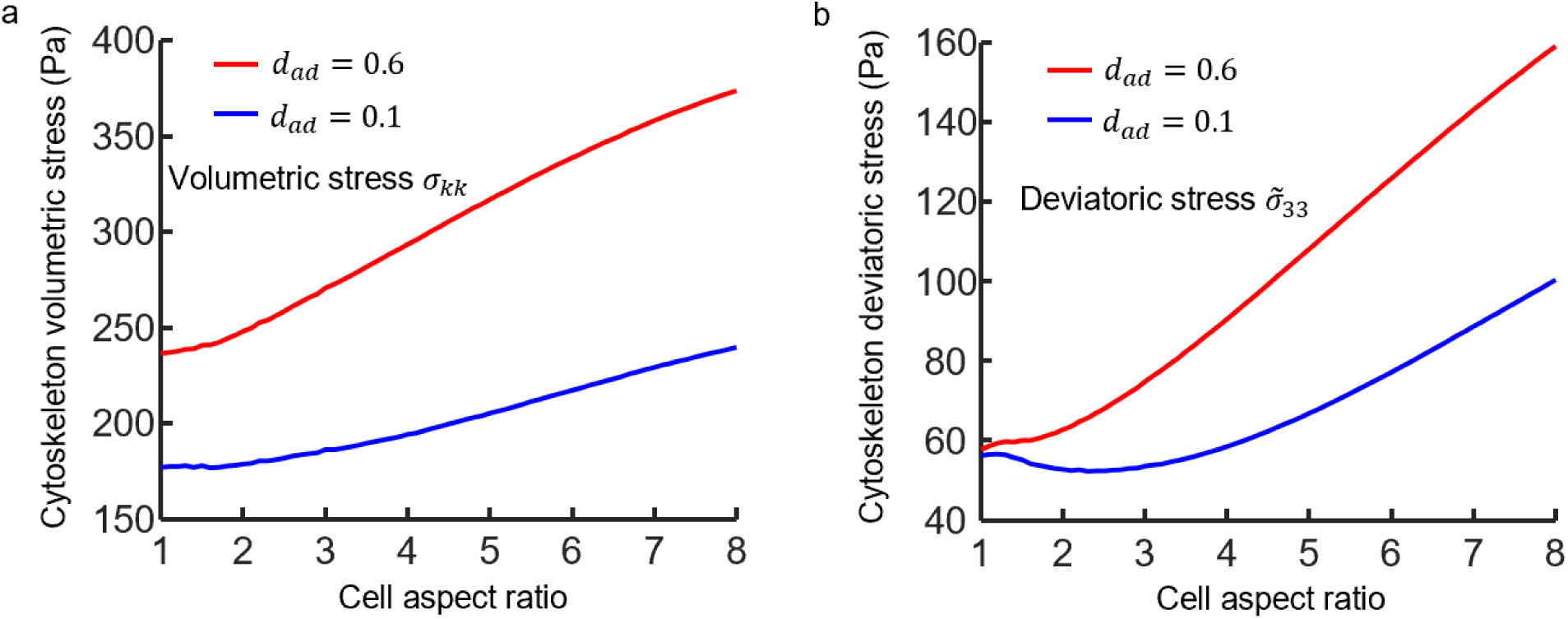
The cytoskeleton (a) volumetric stress and (b) deviatoric stress when the cell integrin-collagen adhesion density is low (*d*_*ad*_ = 0.1) or high (*d*_*ad*_ = 0.6).

**Fig. S5.**
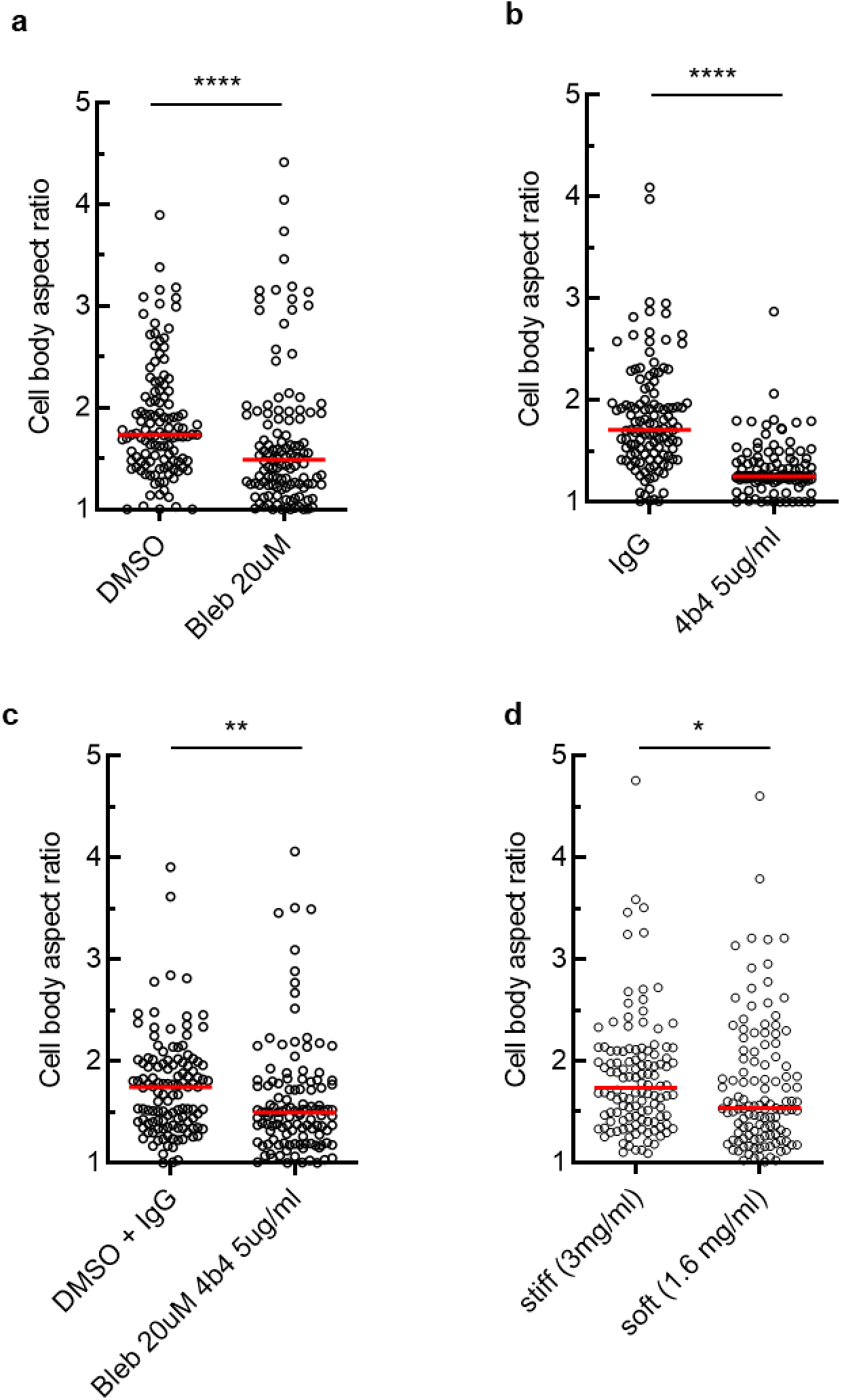
Cell body aspect ratio of HT1080 cells migrating for 20hr in 3mg/ml bovine collagen under treatment (a,b,c) or untreated in different collagen densities (d) as indicated. 40 cells/condition (three independent experiments). Bleb, Blebbistatin. Mann-Whitney U test (****, p-value <0.0001; **, p-value =0.0012; *, p-value =0.0185).

**Fig. S6.**
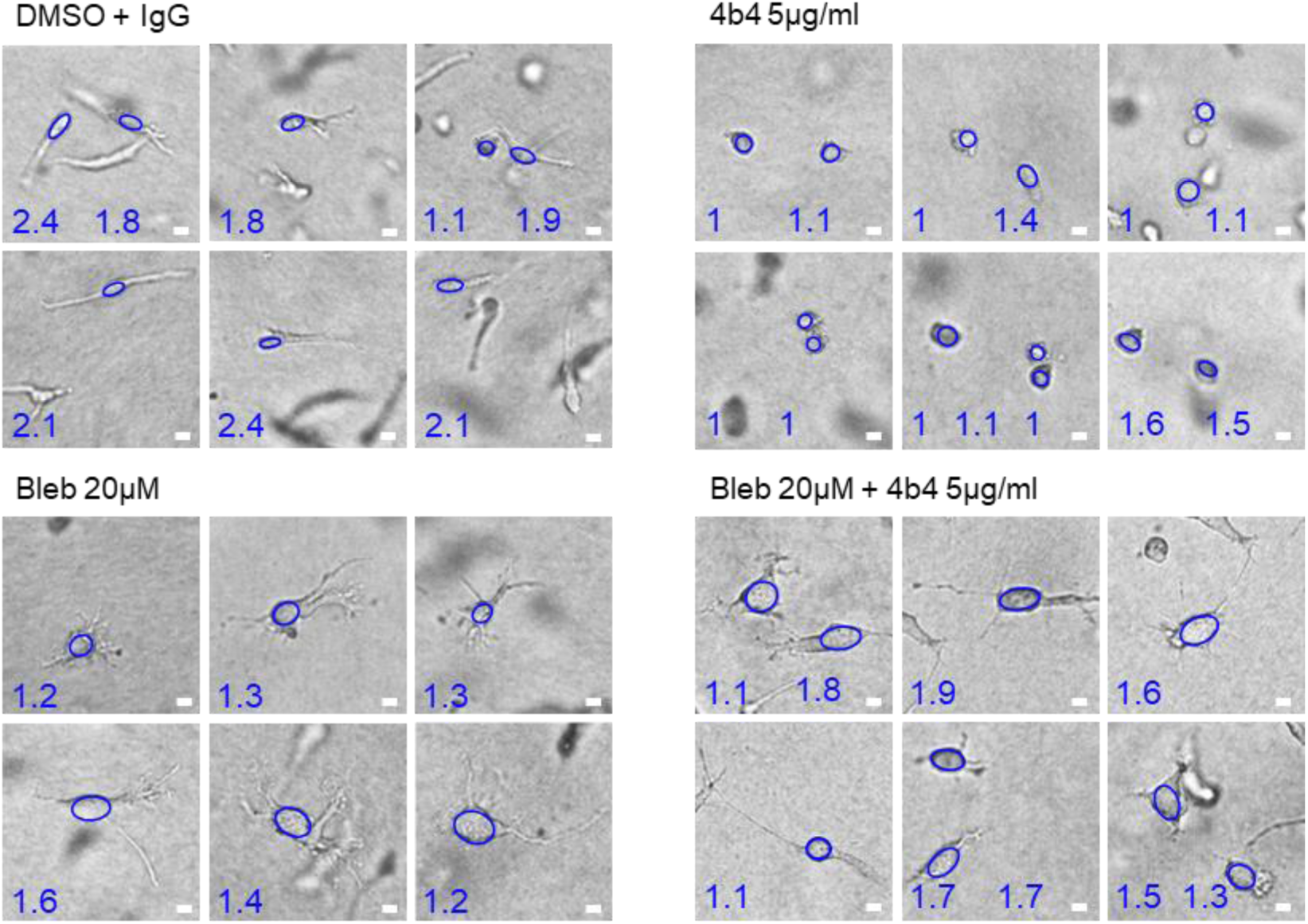
Representative cell morphologies of HT1080 cells migrating for 20hr in 3 mg/ml bovine collagen with or without mono- or –dual-interference of myosin II mediated cell contractility (Bleb, Blebbistatin treatment) and β1 integrin mediated cell-ECM adhesion (4b4 treatment) recorded by bright-field microscopy. Ellipses indicate cell bodies, values represent respective cell body aspect ratios. Scale bars, 10μm.

## Supplementary information

### 1. The steady state of the cell in the matrix

We consider the timescale of cell morphogenesis is significantly longer than the time required for the cell-matrix system to reach a steady state, in which no macroscopic changes occur (e. g. *ϵ*_*ij*_, *ρ*_*ij*_ becomes constant). With this assumption, the free energy of a cell with a given shape is computed based on the steady-state stress/strain fields. Using the energy functions specified above (Eq. 1-8), the true stress in the cell 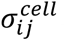, cell-matrix interface 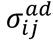 and the matrix 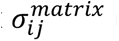 can be written as

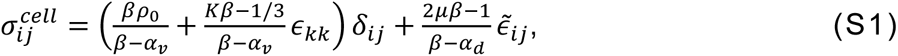

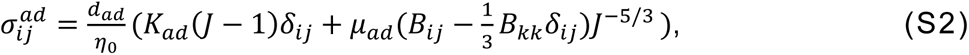

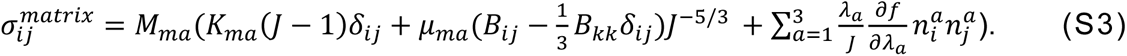

In these equations, *δ*_*ij*_ denotes the Kronecker delta, *n*^*a*^(*a* = 1,2,3) are the unit vectors along the three principal directions, *J* = *λ*_1_*λ*_2_*λ*_3_, and *B*_*ij*_ denotes the left Cauchy-Green tensor. The steady state stress/strain fields are then solved by applying boundary conditions and force-balance equation *∂σ*_*ij*_/*∂x*_*i*_ = 0 (see SI for details). Note that for the sake of simplicity, we ignored the adhesion binding affinity, membrane and cortical tension when calculating the stress in the cell-matrix interface (Eq. 10).

### 2. Constitutive models

#### ECM

The ECM is treated as a fibrous network with constitutive model that we developed recently^13^. Based on the discrete fiber simulations in our previous study, there are two families of fibers in response to tensile mechanical loading: the fibers oriented randomly, and the fibers aligned along the directions of tensile principal stretches. The strain energy density function of the ECM consists of the contributions from both the unaligned and aligned fibers, which can be written as

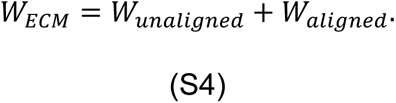

The unaligned fibers display an isotropic mechanical behavior, which can be characterized by the strain energy density function

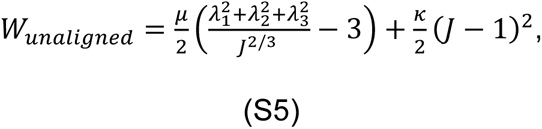

where *λ*_1_, *λ*_2_ and *λ*_3_ are the stretches in the three principal directions. *J* = *λ*_1_*λ*_2_*λ*_3_ denotes the deformation volume ratio of the network. This constitutive relation reduces to linear elastic model at small strains, with shear and bulk moduli given by *μ* and *κ*. The aligned fibers formed due to tensile strains stiffen the network in the alignment directions. With increased loading, the number of the aligned fibers increase, enhancing the network stiffness. This stiffening mechanism of the fibrous network can be captured by

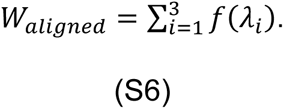

Where *f*(*λ*_*i*_) is defined as

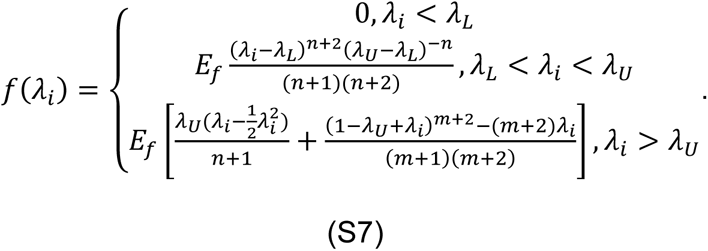

Here *E*_*f*_ is the parameter that characterizes the stiffening due to fiber alignment, and

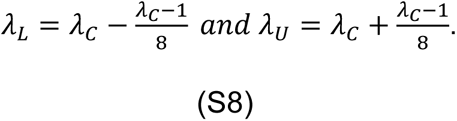

*λ*_*c*_ is the critical strain beyond which the collagen fibers start to align and stiffen.

*W*_*aligned*_ is the strain energy density of the aligned fibers, *E*_*f*_ and *m* are two parameters determining the stiffness and stiffening behavior. Using Eq. S4 – S8, the principal stresses in the ECM can be calculated by

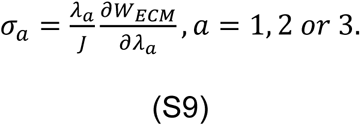

Cell nucleus:

The cell nucleus is treated as a non-linear elastic material, which is characterized by

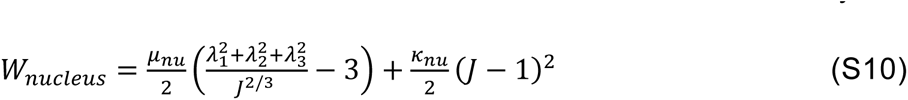

where *λ*_1_, *λ*_2_ and *λ*_3_ are the stretches in the three principal directions. *J* = *λ*_1_*λ*_2_*λ*_3_ denotes the deformation volume ratio of the network. This constitutive relation reduces to linear elastic model at small strains, with shear and bulk moduli given by *μ*_*nu*_ and *κ*_*nu*_.

#### Cell-matrix adhesion

While each of the adhesion connect the cell and the matrix individually, we consider the the collective contribution of all adhesions as a continuum elastic layer. They are treated as an non-linear elastic material, which is characterized by the strain energy density function

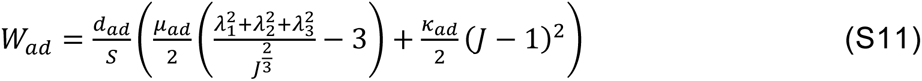

Here in this equation, S is a scale factor that relates the stiffness of the adhesion layer. *λ*_1_, *λ*_2_ and *λ*_3_ are the stretches in the three principal directions. *J* = *λ*_1_*λ*_2_*λ*_3_ denotes the deformation volume ratio of the network. This constitutive relation reduces to linear elastic model at small strains, with shear and bulk moduli given by *μ*_*ad*_ and *κ*_*ad*_. The strain energy of the adhesion is then calculated by integrating *W*_*ad*_ over the volume of the adhesion layer *V*_*ad*_

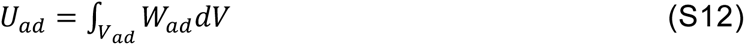

### 3. Degradability of the matrix

It is important to note that our model assumes cells remodel the surrounding fibrous matrix, by secreting matrix metalloproteinase (MMP) to degrade the network. With this assumption, there is no matrix stress induced by a change in cell shape. Conclusions may therefore vary for cells in a non-degradable matrix, or if cell MMP expression be inhibited. In such cases the cells are restricted by the matrix fiber architecture. When the matrix pore size is small, cells are confined within the matrix pores and lose the ability to migrate^18^, and cell shape is constrained by pore shape^27^. For a fully non-degradable matrix, any change in cell shape can lead to high matrix stresses, increasing *F*_*matrix*_ significantly and penalizing shape change. With an increase in matrix degradability, the regulatory role of *F*_*cell*_ and Γ on shape will gradually be more significant. This effect has been observed in experiments that compare cell morphology in degradable and non-degradable gels. A study by Anderson et al.^27^ revealed that human mesenchymal stem cells (hMSCs) spread into spindle-shaped morphologies in MMP-degradable gels. However, in less-degradable gels cell morphology was more rounded.

